# Stratified time-course gene preselection shows a pre-diagnostic transcriptomic signal for metastasis in blood cells: a proof of concept from the NOWAC study

**DOI:** 10.1101/141325

**Authors:** Einar Holsbø, Vittorio Perduca, Lars Ailo Bongo, Eiliv Lund, Etienne Birmelé

## Abstract

We investigate whether there is information in gene expression levels in blood that predicts breast cancer metastasis. Our data comes from the NOWAC epidemiological cohort study where blood samples were provided at enrollment. This could be anywhere from years to weeks before any cancer diagnosis. When and if a cancer is diagnosed, it could be so in different ways: at a screening, between screenings, or in the clinic, outside of the screening program. To build predictive models we propose that variable selection should include followup time and stratify by detection method. We show by simulations that this improves the probability of selecting relevant predictor genes. We also demonstrate that it leads to improved predictions and more stable gene signatures in our data. There is some indication that blood gene expression levels hold predictive information about metastasis. With further development such information could be used for early detection of metastatic potential and as such aid in cancer treatment.

## I. Introduction

About one in ten women will at some point develop breast cancer. About 25% have an aggressive cancer at the time of diagnosis, with metastatic spread to axillary lymph nodes. ^1^ Spread is detected by a sentinel node biopsy: a surgical procedure to check the lymph nodes closest to the cancer site for metastasized cancer. A cancer that has developed to the point of metastasis is much more dangerous than a local one. The absence or presence of metastatic spread largely determines the patient’s survival. Early detection is hence very important in terms of reducing cancer mortality. Were we able to detect signs of metastasis or metastatic potential by a blood sample, perhaps in a screening setting, we could conceivably start treatment earlier and treat the cancer before the onset of large, deadly metastasized tumors.

Several recent articles develop this idea of *liquid biopsies* [1]. Different relevant signals appear in blood for already diagnosed breast cancer. For instance: circulating tumor cells [2], circulating tumor DNA [3], serum microRNA [4], or tumor-educated platelets [5]. A recent review in *Cancer and Metastasis Reviews* [6] lists liquid biopsies and large data analysis tools as important challenges in metastatic breast cancer research.

Norwegian Women and Cancer (NOWAC) [7] is a prospective study that includes blood samples. A subset of these have been processed on microarrays to provide gene expression measurements in the form of mRNA abundance (transcriptomics). Any information contained in blood gene expression will be of a systemic nature. Since cancer grows over time, so presumably does any systemic response increase over time accordingly. This would mean that blood samples provided long before a cancer has grown enough to be detectable and diagnosable necessarily holds less information than a more recent blood sample. Hence prospective blood samples provide gene expression trajectories over time. Such trajectories should diverge between cases and controls as the tumor grows. Lund et al. [8] show a significant difference in trajectories for groups of genes. In this paper we aim to show that we can go further and find predictive information about metastatic spread to the closest lymph nodes (the sentinel nodes).

Making predictions from gene expression is a high-dimensional problem. There are about ten thousand potential predictor genes. In such a setting it is very easy to find noise that looks like a signal. The hope is usually to uncover lower-dimensional structures in the data. For instance, we expect genes to work together in pathways. We do not expect all genes to be relevant in all processes. The analysis of high-dimensional data is an active research area of statistics and machine learning [9]. The common methods for discovering low-dimensional structure are projection approaches like PLS-methods [10] and variable selection.

Variable selection approaches highlight the most discriminative variables. A set of selected variables is simpler to interpret than a projection. There is a variety of variable selection schemes; Fan and Lv [11] provide a review. Working with gene expression we can rank genes for example based on genewise t-tests for differential expression. The top *k* of these provide a lower-dimensional space where we can apply any prediction method. Haury et al [12] show that such a ranking coupled with a simple classifier method compares favorably to more sophisticated methods. There are also integrated methods that do simultaneous selection and model fitting. A popular choice is the penalized maximum likelihood family of generalized linear models, which optimize the likelihood plus a penalty term that encourages sparse solutions. These include the popular lasso and elastic net methods [13].

The chosen set of predictors is often unstable in that they change with perturbations to methodology or data set. Ein-Dor et al. [14] examined the effect of using different subsets of the same data to choose a predictive gene set. They show that predictor gene sets depend strongly on the subset of patients used for analysis. Stability criteria make a complimentary feature to predictive power. These can be integrated in the model selection, as the stability selection for penalized regression [15], or they can be used for posteriori evaluation [12].

In this article we examine some feature selection schemes for metastasis prediction with blood samples from a prospective study. We use gene expression data from 88 case–control pairs from the NOWAC study. The blood samples were provided 6–358 days before diagnosis. We propose variable selection based on a gene’s prediagnostic progression over time and the manner in which the cancer was diagnosed. We also provide a means to simulate prospective gene expression data, extending a previous method.

For both variable selection and prediction we use simple, well-established methods. We do not attempt to provide a survey and comparison of advanced techniques: the size of the data only permits approximate insights. For this same reason we do not enumerate any predictive gene set—in fact we shall see that such gene sets are quite unstable. Biologically we see some evidence of a signal predictive of metastasis already present before diagnosis in these data. This gives some hope for the pursuit of liquid sentinel node biopsies for diagnosis or crucial early detection. Statistically we see that the use of followup time and especially stratification is important to find this information.

## II. Material and methods

### A. Data

We analyze 88 pairs of cases with breast cancer diagnoses and age-matched healthy controls from the NOWAC Postgenome cohort. Dumeaux et al. describe the NOWAC study in detail [7]. In brief, women in a certain age group received an invitation to participate by random draw from the Norwegian National Registry. The women who chose to participate filled out a questionnaire and provided a blood sample. Over the years the Cancer Registry of Norway provided followup information on cancer diagnoses and lymph node status. The women in this particular data set received a breast cancer diagnosis at most one year after providing a blood sample.

The NTNU genomics core facility processed the blood samples with Illumina microarray chips of either the HumanWG-6 v. 3 or the HumanHT-12 v. 4 type. To keep case–control pair as comparable as possible, the pair is intact throughout processing pipeline. This means that they are processed on the same day by the same person and lie next to one another physically on the microarray chip. All NOWAC data sets go through a standardized technical quality control [16]. We have removed low-signal and low-quality probes. Finally we have normalized the data by quantile normalization before analysis. The preprocessing for these particular data is described in detail by Lund et al. [8].

The end-result is a 88 × 12404 **fold change matrix**, *X*, on the log_2_ scale. For each gene, *g*, and each case–control pair, *i*, we have the measurement 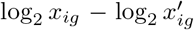. Here *x_ig_* is the *g* expression level for the *i*th case, and 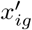 is the corresponding control. For each case we have the number of days between the blood sample and the cancer diagnosis. We will call this the **followup** time, or simply the followup of a case. Although followup introduces a time aspect, these are not time series data. Each observation is a different woman, so there should be no autocorrelation to speak of, and followup time is random. The variable **detection stratum** takes one of the following values:

- *Screening* denotes a cancer that was detected in the regular screening program.
- *Interval* denotes a cancer that was detected between two screening sessions. The interval between screenings is two years.
- *Clinical* denotes a cancer that was detected outside of the screening program. These women either never took part in the screening program, or did not attended a screening in at least two years.

The response variable, **metastasis** (∈ {0,1}), indicates whether a sentinel node biopsy showed evidence of metastasis. We sometimes refer to this as spread. Table I shows the incidence of metastasis in the different strata. There is a certain heterogeneity. We conjecture that an interval cancer grows rapidly, having appeared in the two years between screens, and that a clinical cancers has been growing for a long time before becoming problematic enough that the patient suspects something herself.

**Table I.**
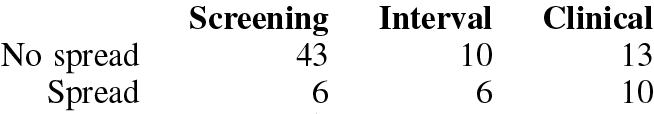
Incidence of metastasis across detection strata. There is noticeable between-stratum variation. The incidence is much lower in the screening stratum.

### B. Variable selection

We investigate four ways to rank genes, which we describe briefly in this section. The methods all assess differential expression between groups in some way. We propose the first—ANOVA—to take into account a hypothetical functional relationship between gene expression and time. The other three—SAM, t-tests, and LIMMA moderated t-tests—are well-established methods for ranking genes and assessing differential expression.

#### 1) ANOVA

We hypothesize that the expression of genes that are relevant to the cancer process diverges over time. To detect this behavior we regress fold change, *e*, on time, *t*, and metastasis, *M*, in the following model:

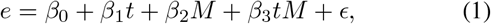

where *ϵ* is iid noise. We refer to this as **ANOVA-f** below.

We suspect that different genes may be relevant in different detection strata. We model this by expanding equation 1 to include an interaction with stratum, *S*:

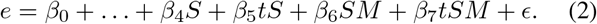

We refer to this as **ANOVA-fs** below.

Finally we entertain the possibility that followup is not important and that stratum alone is of interest. This yields the model

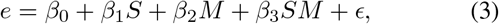

which we refer to as **ANOVA-s** below. Note the abuse of notation in Equations 2 and 3; *S* has three levels and will be coded as a dummy variable.

We rank genes by the *F*-statistic obtained under the null hypothesis that the model in Equation 1, 2, or 3 is no better than the intercept-only model, *e* = *β*_0_ + *ϵ*. Ignoring both stratum and followup is equivalent to a regular t-test for metastasized vs not as in Section II-B2 below.

#### 2) t-test

We rank genes by Welch’s two-sample t-statistic [17] between metastasized and non-metastasized cases (**t-test** below). This is complementary to the three methods above, as regressing on a single binary grouping variable can be used as a test for difference in means.

#### 3) SAM

The Significance Analysis of Microarrays (**SAM**) procedure of Tusher, Tibshirani and Chu [18] defines the relative difference in gene expression for the *i*th gene as:

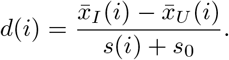

Here 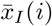 and 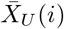 are the average expression levels of gene *i* in the two states *I* and *U* (metastasized or not), *s*(*i*) is the pooled standard deviation estimate in the two states, and finally *s*_0_ is a small positive constant added to all genes to make the variance of *d_i_* independent of gene expression level. We rank genes by *d*(*i*).

#### 4) LIMMA t-test

Smyth’s Linear Models for Microarray Data (LIMMA) is a general empirical Bayes framework for assessing differential expression [19]. The LIMMA moderated t-statistic, 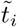, is similar to the SAM *d*(*i*) in that it modifies the denominator of a regular t-statistic. In this case we have that for the *i*th gene,

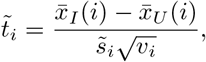

where *v_i_* is a factor that has to do with the variance of 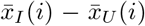. The standard deviation estimate 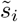 has been shrunk by empirical Bayes methods toward the average standard deviation across all genes. We refer to this as **LIMMA-t** below.

### C. Prediction

Having ranked genes and chosen the top *k* as predictors, we use these in the following logistic regression model for the probability, *p*, of metastasis:

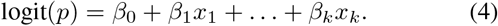

This model uses only gene expression levels regardless of whether stratum (or time) was used in selecting the predictors. This model can be used in a screening setting (where the cancer has not yet happened and hence we do not have information about its detection). Since followup time is a result of these data coming from a cohort study, it could not be used in a realistic predictive model.

Considering Table I it is likely that detection type is informative of the probability of metastasis. Conceivably a predictive model could be used at time of diagnosis where this information is available. Our model for such a setting is simply Equation 4 with an extra interaction with stratum, much like in the variable selection in Section II-B1:

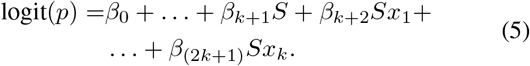

We estimate models 4 and 5 by Bayesian generalized linear models with a weakly informative prior from Gelman et al. [20]. This is more for convenience than from a particular wish to do Bayesian modeling: when selecting the *k* “best” predictors out of thousands of candidates it is quite likely to find some where the metastasis and non-metastasis points are linearly separable (ie. their respective convex hulls are disjoint). In such a setting, the classical iteratively reweighted least squares optimization does not converge. Predictors selected for some function of their effect size would likely regress toward the mean in new data and some amount of shrinkage is prudent. The standard prior of Gelman et al. provides a sensible and convenient regularization without a need for parameter tuning.

### D. Baseline

We compare predictive performance against two naive and two more sophisticated baselines. The more sophisticated baselines use all genes without prior ranking and selection.

The first naive model we consider is the “random guess” intercept-only model logit(*p*) = *β*_0_. Second we compare against using the stratum information, logit(*p*) = *β*_0_ + *β*_1_*S*, corresponding to making a recommendation based only on the manner in which the cancer was detected.

The other two baselines are penalized logistic regression models. These models take the same form as Equation 4, using all predictors rather than the top *k*, but maximize the likelihood subject to the constraint that 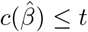. Ie. the magnitude *c*(·) of the coefficients *β_i_* must not exceed some threshold *t*. We investigate ridge penalty, 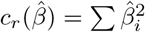 and Tibshirani’s lasso penalty, 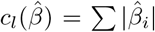 [21]. These are well-known models. The lasso provides an end-to-end solution that does variable selection and model fitting in one go. Using a ridge penalty simply uses all predictors but shrinks coefficients toward zero quadratically in their magnitude. Both of these methods require the selection of *t*; we choose this by cross validation.

### E. Metrics

We evaluate models by three criteria. **Brier score** [22] is the mean squared error,

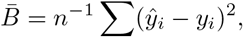

between the probability that was predicted by the model, *ŷ*, and the known outcomes, *y*. **Concordance probability** is the probability of ranking (in terms of probability) a randomly chosen positive higher than a randomly chosen negative (ie a metastatic vs a non-metastatic case, respectively). This is equivalent to area under the receiver operating characteristic curve (AUC), and is proportional to the Mann-Whitney Wilcoxon U statistic [23]. Finally, **stability** is the probability of recovering the same predictor genes between different realizations of the modeling procedure. We follow Haury et al. [12] and measure this by the Jaccard index, 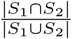, where *S*_1_ and *S*_2_ are two sets of predictor genes.

### F. Validation

#### 1) Optimism bootstrap

For estimation of concordance probability and Brier score we take the optimism-corrected bootstrap approach described amongst other places in Efron and Gong [24]. This has the advantage of using all of the data in estimating model performance opposed to data splitting procedures such as out-of-bootstrap or *k*-fold cross-validation where only a portion of the data is used to fit the model.

The apparent score (or training score), 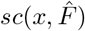, is the expected score of a model fit to the sample, *x*, w.r.t. the empirical distribution of this sample, 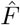. This is necessarily an overoptimistic estimate. To correct for this, we estimate the expected overoptimism, *ω*, by the bootstrap:

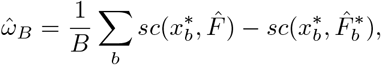

where 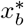 is the *b*th bootstrap sample and 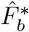 is the empirical distribution function of the same. Hence 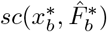 is the apparent score of the *b*th bootstrapped model, and 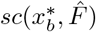 is the expected score of the same model w.r.t. the empirical distribution of the original sample. The optimism-corrected expected score of our model becomes

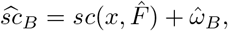

which is a bias correction of the apparent score.

It does not make sense to estimate stability by optimism correction. Let *S*(*X*) be the gene set selected in the original data and 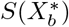 be the gene set selected in the *b*th bootstrap sample. The bootstrap estimate of expected stability is then

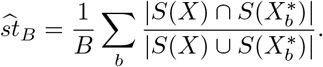

#### 2) Standard errors

We measure uncertainty in the bootstrap estimates by the jackknife-after-bootstrap procedure. The jackknife estimate of standard error for any statistic 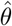 is

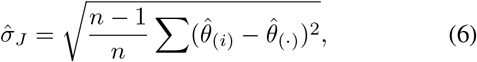

where 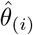 is the statistic computed with the *i*-th sample removed, and 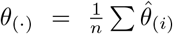. In principle the bootstrap procedure has to be repeated for each 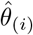. But there is a computational shortcut here due to the fact that a bootstrap sample drawn with replacement from *x*_1_, …, *x*_*i*−1_, *x*_*i*+1_, …*x_n_* has the same distribution as a bootstrap sample drawn from *x*_1_, …*x_n_* in which *x_i_* does not appear (the *jackknife-after-bootstrap lemma* in Efron and Tibshirani [25]).

### G. Simulation study

To simulate a time-course, stratified gene expression data we extend the scheme of Dembeléeé [26]. We wish to compare ranking methods in a simple setting, do not consider metastasized cases vs non-metastasized cases and examine only a general case vs control scheme.

Briefly, Dembélé’s scheme is to model the the log_2_ gene expression levels *x* for gene *i* as

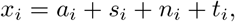

where *x_i_* is a vector of the observed values of gene *i* for all subjects. Vector *a_i_* contains baseline expression values, while *s_i_* defines offsets for differential expression. The offset is zero for controls and genes that are not differentially expressed. The vector *n_i_* contains independent noise and the vector *t_i_*, unused, models technical errors.

We modify the gene expression component *a_i_* + *s_i_* so that

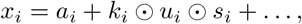

where ⊙ denotes element-wise multiplication. The factor *k_i_* introduces a decay in effect size for longer followup time. We sample a followup-time vector *t_i_* ~ *U*[0; 1]^*n*^ and set *k_i_* = 1 − *t_i_*. This shrinks differential expression toward zero proportinally to followup time. The factor *u_i_* introduces a stratum effect. We first draw a stratum indicator *r_ij_* ~ Bernoulli(.5), and then let *u_ij_* = .5 if *r_ij_* = 1 and *u_ij_* = 1 otherwise. This gives us the stratum effect vector *u_i_* = [*u_ij_*]_*j*=1…*n*_. The expected differential expression in stratum 1 is half of that that in stratum 0.

Dembélé’s method is implemented in the R-package `madsim`. It allows one to specify an empirical distribution to sample from. We use 30 000 random gene expression values from our real data as a seed to the algorithm. There are several parameters to the method. Important to us are 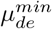 and λ_2_. Differential expression is realized 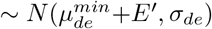, where *E*′ ~ *exp*(λ_2_). Hence 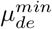 is the smallest possible average differential expression, and λ_2_ a parameter that adjusts how often average differential expression departs noticeably from this minimum. We set 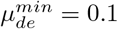 and *σ_de_* = 0:025, to make the smallest possible effect quite small and stable. The noise component is *n_i_* ~ *N*(0; σ*_n_*). We set σ*_n_* = 0:2 to roughly match the variance we see in our own data. This gives us a signal-to-noise ratio of .5 in the worst case. The expected differential expression in this model is 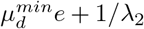. We let λ_2_ be an adjustable parameter to aid our investigations.

We make our modified simulation scheme available as an R package at https://github.com/3inar/pmadsim. This is a fork of `madsim` and shows all modifications we have made to the original package. It provides a function that generates data according to the above specifications.

## III. Results

This section presents our results. First those from our simulation, then those from predicting metastasis in the NOWAC data.

### A. Simulations

We generate 10 000 gene expression measurements for 33 cases and 55 controls. This roughly matches our data set, which has about 12 000 expression measurements for 33 metastasized cases and 55 non-metastasized cases. We set the probability of differential expression to 0:01 for around 100 differentially expressed genes.

We let λ_2_ range over a set of values so that the expected value *β* = 1/λ_2_ of the corresponding exponential is a series of ten evenly spaced numbers between 0:01 and 2 for an expected effect size from 0:02 to 2:01. For each λ_2_ and each selection method we do 1 000 simulations.

We measure the probability of recovering the 100 differentially expressed genes when picking the top 100 genes as ranked by the different selection methods. Figure 1 shows the results in the presence of a simulated followup decay effect. The differences between methods is slight, but ANOVA-f yields a slightly higher selection probability for all effect sizes (smaller effect sizes omitted in the figure). Figure 2 shows the results in the presence of both a followup and a stratum effect. Again the differences are slight. ANOVA-f still does best, outperforming the correct model of ANOVA-fs. This is likely because ANOVA-fs must estimate three extra parameters per gene with the same small amount of data.

**Figure 1.**
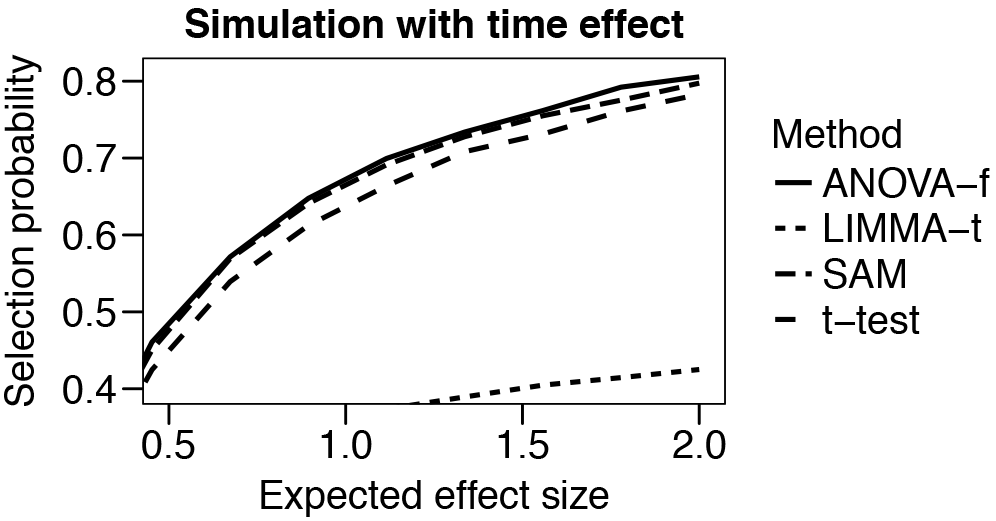
Probability of selecting a truly differentially expressed gene when there is a linear time effect in our simulation. ANOVA-f, SAM and t-tests behave similarly, but accounting for time with ANOVA-f yields a slightly higher selection probability.

**Figure 2.**
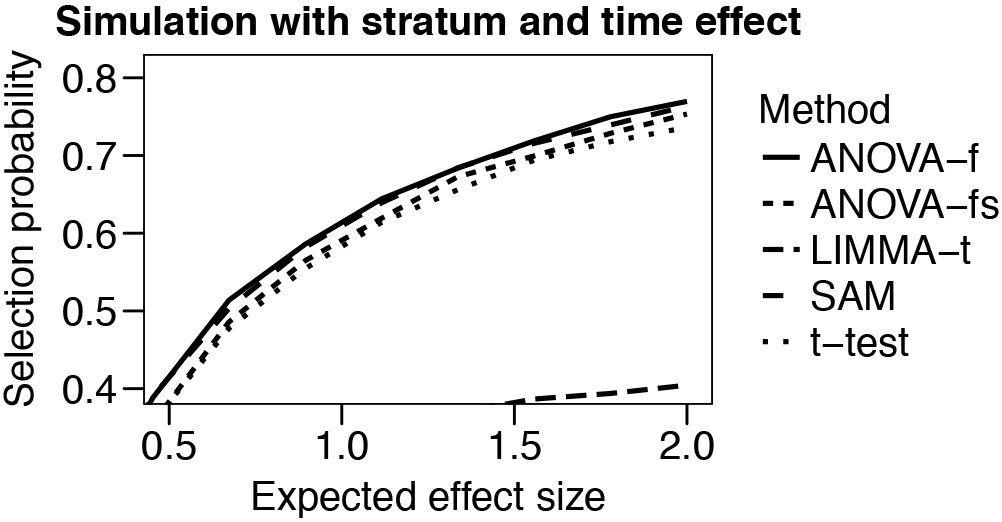
Probability of selecting a truly differentially expressed gene when there is a linear time effect and a stratum in our simulation. Both ANOVA methods, SAM and t-tests behave similarly. Accounting for time with ANOVA-f yields a slightly higher selection probability. However accounting for stratum with ANOVA-fs does not in this case increase the selection probability further.

### B. NOWAC metastasis prediction

Below all bootstrapped results are based on 2 500 resamples. For all ranking methods we choose the top ten genes and use them as predictors for the models described in Equations 4 and 5. This is based on the folk wisdom to have around ten observations per estimated parameter in the regression model. In practice we end up with fewer observations per parameter, especially for model 5, so the models are slightly over-parameterized. This likely contributes to uncertainty in our results.

Tables II and III show the Brier score and AUC for predictions based on the different selection schemes we investigate. Models 4 and 5 refer to Equations 4 and 5 in Section II-B1. That is, respectively, the “screening” prediction using only gene expression in the model, and the “at diagnosis” prediction that uses the additional information of detection stratum.

**Table II.**
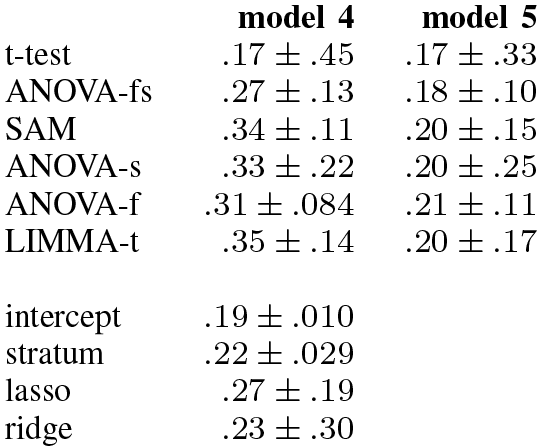
Brier scores presented as point estimate plus/minus two standard errors. Measures error in forecast probability: lower is better. Model number refers to the equations in Section ii-c. Model 5 includes stratum as a predictor. Below the break are the four baseline models.

Brier score is a measure of error: the lower the better. Table II shows Brier score as point estimate plus/minus two standard errors in decreasing order by Model 5 point estimate. The results do not suggest any simple interpretation, but it is noteworthy that the intercept-only model is among the best calibrated. The uncertainty is large enough that is difficult to say that any selection method is better than any other. It is clear that the interaction with detection method in model 5 improves calibration for all models. There is also lower uncertainty in the ANOVA-f/fs models.

AUC or concordance probability is a measure of a model’s ability to discriminate between outcomes: the higher the better. Brier score alone does not provide full information about predictive performance; the intercept only model is well calibrated but cannot be used for prediction at all. Random guess (or forecasting a constant for every observation) yields AUC of .5; perfect discrimination yields AUC of unity. Table III shows AUC as point estimate plus/minus two standard errors in decreasing order by model 5. Again the clearest signal is that the added information from detection method is very important. Point estimates improve markedly and standard errors generally decrease. Also here does use of stratification and followup time in preselection reduce uncertainty.

**Table III.** AUC presented as point estimate plus/minus two standard errors. Measures the probability of forecasting a higher probability of metastasis for a randomly chosen metastasis case than for a randomly chosen non-metastasis case: higher is better. Model number refers to the equations in Section ii-c. Model 5 includes stratum as a predictor. Below the break are the four baseline models.

|  | model 4 | model 5 |
| --- | --- | --- |
| LIMMA-t | .44 ± .30 | .76 ± .20 |
| SAM | .46 ± .26 | .75 ± .24 |
| ANOVA-fs | .51 ± .29 | .75 ± .16 |
| ANOVA-s | .41 ± .57 | .75 ± .38 |
| t-test | .65 ± 1.5 | .74 ± .71 |
| ANOVA-f | .44 ± .25 | .72 ± .21 |
| intercept | .5 |  |
| stratum | .49 ± .055 |  |
| lasso | .36 ± 1.4 |  |
| ridge | .81 ± 3.3 |  |

The ridge regression baseline performance has a very good AUC point estimate, but the standard error is very large. Too large: it is a theorem that the upper bound on standard deviation in a variable ∈ [0, 1] is 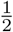. This says something about the imperfection of the jackknife as an estimator of standard error. The blame lies at least in part with the correctional factor 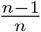 in Equation 6, which was originally defined heuristically. Since it is difficult to suggest a sensible alternative, we choose to live with this.

The collected results for model 5 suggest some reason for optimism. Due to the size of the standard errors we must necessarily be uncertain about even the first significant digit of our point estimates. But even accounting for uncertainty there seems to be predictive information better than random guess. As in the simulations, there is not too much difference between the different methods, perhaps apart from the simple t-test, for which we observe much variance. Note that both SAM and LIMMA are flexible frameworks and we could have accounted for stratum and followup in either. Our comparison is between using this information and various ways of not using it, and there is no reason to believe that either framework should perform poorly if we were to use more refined models there.

Table IV shows the predictor set stability as point estimate plus/minus two standard errors. Stability is in general very low, and the standard errors suggest that there is even some uncertainty to the order of magnitude of the point estimates. A possible interpretation is that the correlation between genes is such that many different genes hold similar information. It is at least clear that we need much more data if we want to find a stable set of predictor genes. If we take the point estimates at face value, Table IV reflects the fact that we see lower uncertainty using ANOVA-f/fs in Tables II and III.

**Table IV.**
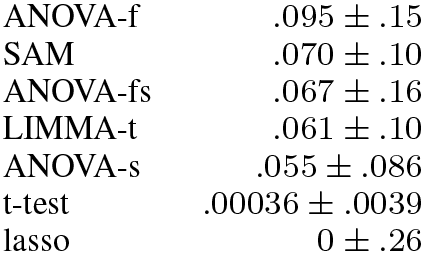
Stability as point estimate plus/minus two standard errors. Stability is an estimate of the probability of recovering the same gene set with different realizations of a modeling procedure. A larger stability provides more certain biological interpretation. The lasso is the only baseline method included here as it is the only one that does variable selection.

|  |  |
| --- | --- |
| ANOVA-f | .095 ± .15 |
| SAM | .070 ± .10 |
| ANOVA-fs | .067 ± .16 |
| LIMMA-t | .061 ± .10 |
| ANOVA-s | .055 ± .086 |
| t-test | .00036 ± .0039 |
| lasso | 0 ± .26 |

## IV. Conclusion

We have found some indication that predicts breast cancer metastasis could be predicted from blood transcriptomic measurements. The information about how a cancer was detected is very important in this respect. This suggests separate mechanisms at work in the different strata and that perhaps these three settings should be explored separately. These results are a long way from practical application, but the potential impact of early detection of metastasis is great. Larger studies should be conducted to provide reliable evidence, especially if there is to be any hope for identifying a stable gene signature set for metastasis.

## Footnotes

1 http://oncolex.org/Breast-cancer/

